# Single-cell proteomics of pancreatic islet cells reveals type 1 diabetes and donor-specific features

**DOI:** 10.1101/2025.09.18.677038

**Authors:** Pierre Sabatier, Louise Granlund, Maico Lechner, Sergey Rodin, Karl-Henrik Grinnemo, Olle Korsgren, Jesper V. Olsen

## Abstract

Type 1 diabetes mellitus (T1DM) is the most common severe chronic disease in children and adolescents and requires life-long exogenous insulin treatment. Investigating key differences between the pancreas of healthy and T1DM patients is key to help uncover new treatments. Here we performed a single-cell proteomic (SCP) analysis of purified pancreatic islets from several healthy and T1DM donors at a depth of > 4000 protein quantified. We employed tools designed for single-cell mRNA sequencing (scRNAseq) for cell type annotation highlighting discrepancies between mRNA and protein markers. Subsequently, we defined a subset of markers useful for future proteomics analyses of pancreatic islets. This marker list was employed to refine the annotations of our combined dataset, which we exploited with a machine learning approach to successfully transfer annotations into a new islet dataset. Lastly, we showcase the applicability and value of SCP in a clinical context by providing novel insights into cell type-specific differences between healthy and T1DM islets as well as between healthy islets of different donors. Our study is the first SCP analysis of pancreatic islets from multiple donors and T1DM islets, performed at this level of analytical depth. This provides methodological and biological insight as well as a reference dataset for future SCP studies and for any researcher interested in T1DM and in pancreatic islet physiology.

## Introduction

The endocrine pancreas comprises several distinct cell types, organized into islets of Langerhans. The islets consist of at least five different endocrine cell types: alpha, beta, delta, pancreatic polypeptide and epsilon cells, secreting insulin, glucagon, somatostatin, pancreatic polypeptide and ghrelin, respectively^1,2^. The vast majority of the pancreatic cells are ductal and acinar cells, producing and delivering exocrine enzymes into the duodenum, in total the endocrine islet cells only constitute about 1-2 % of the total mass of pancreas^3^. Nevertheless, these cells are essential in maintaining blood glucose homeostasis^4^ and play a central role in the development of diabetes, which is marked by the loss of functional beta cells. In T1DM, the majority of the insulin-producing beta cells are lost, however, in almost all subjects affected a small number of beta cells remain decades after diagnosis^5^. T1DM is a chronic, incurable condition, leaving patients dependent on exogenous insulin—the same therapy introduced more than a century ago—despite decades of intensive research. In recent years, efforts to collect and share diabetic pancreatic tissue have increased, yet samples from T1DM cases remain extremely limited.

Single-cell approaches can reveal the diversity and specific protein expression patterns of the distinct cell types composing organs such as the pancreas, which bulk analyses would miss. This is especially important for characterizing and understanding both healthy and T1DM islets, which are highly heterogeneous. Understanding the protein landscape of individual islet cells can help identify new biomarkers for early diagnosis, monitor disease progression, and develop targeted therapies or cell replacement strategies. Single-cell proteomics (SCP) can identify protein alterations in specific cell types during disease progression, thereby clarifying the molecular mechanisms driving beta cell dysfunction and loss. Recent years have witnessed remarkable progress in both the sensitivity and throughput of SCP, driven by innovations in sample preparation workflows and mass spectrometry (MS) acquisition strategies^6–17^. State-of-the-art SCP workflows such as the proteoCHIP EVO 96 and the Chip-Tip workflow, enables the identification of >5,000 proteins from individual cells in a high-throughput and nearly loss-less manner^9,18^. With these technological advances, single-cell proteomics now offers coverage almost on par with scRNAseq albeit at a much lower throughput. Since proteins are regulating and performing most cellular functions and mRNA levels often correlate poorly with their corresponding protein levels, SCP has high potential for functional pre-clinical studies and has, in theory, better chances at finding new protein biomarkers or target candidates than scRNAseq^19^. In this study, we had the rare opportunity to perform single-cell proteomic analysis on freshly isolated pancreatic islets from organ donors with and without T1DM.

## Results

To obtain pancreatic islets of the highest quality, pancreases from heart-beating organ donors, treated according to organ transplantation protocols, were procured through the Nordic Network for Clinical Islet Transplantation (https://nordicislets.medscinet.com/en.aspx). Throughout the article the islets from healthy donors and patients suffering from T1DM will be refer to as healthy and T1DM donor islets, bearing in mind that while the pancreas of T1DM patient were processed through the same transplantation protocols as healthy islets, they are not meant for transplantation. The islets were isolated according to clinical routine, and the isolated islets were kept under optimal culture conditions (see Methods). The culture times were kept as short as possible (between 2-6 days) before the islets were processed for SCP.

The healthy and T1DM pancreatic islets were dissociated into single cell suspensions and processed through our single-cell proteomics pipeline^9,20^ (Figure 1A). Single cells were sorted and prepared using the cellenONE platform with protein extraction and digestion in the proteoCHIP Evo96. Resulting single cell peptide mixtures were desalted and concentrated on Evotips and subsequently analyzed using the Whisper Zoom gradients on Evosep One liquid chromatography (LC) system connected to an Orbitrap Astral mass spectrometer operated in narrow-window data-independent acquisition (nDIA)^21^. Initially, we processed and analyzed pancreatic islets from three donors, namely healthy 1, healthy 2 and T1DM 1. The purity of the islet extraction was assessed by microscopy, staining for endocrine and exocrine cells (unwanted cells) (Figure 1B). The purity was estimated to be 90%, 89% and 63% for the extraction of islets from healthy 1, healthy 2 and T1DM 1 donors, respectively. Single cells with less than 500 proteins quantified were filtered out resulting in 756 single cell proteomes of high quality. The SCP analysis yielded more than 3000 proteins quantified on average in healthy 1 and healthy 2, while slightly more than 1000 proteins in T1DM 1 (Figure 1C). For best results, the SCP sample preparation has to be performed the same week as the pancreatic islet isolation and since it was impossible to predict when fresh donor material would be available, particularly for T1DM, technical batch effects were observed. This was reflected in protein coverage, which differed depending on sample preparation and analytical performance. The quality of the isolation also had impact on the number of proteins quantified. For instance, islet isolation from donors with T1DM presents several challenges, often resulting in reduced islet purity and lower total islet equivalents (IEQ). These limitations are attributed to multiple factors. These limitations arise from several factors. T1DM donors generally have smaller pancreases, resulting in lower islet yields^22,23^. Elevated HbA1c levels are also negatively correlated with total IEQ, further reducing isolation efficiency. In addition, islets from T1DM donors often display no or weaker dithizone staining, complicating image-based quantification and analysis^24^.

**Figure 1.**
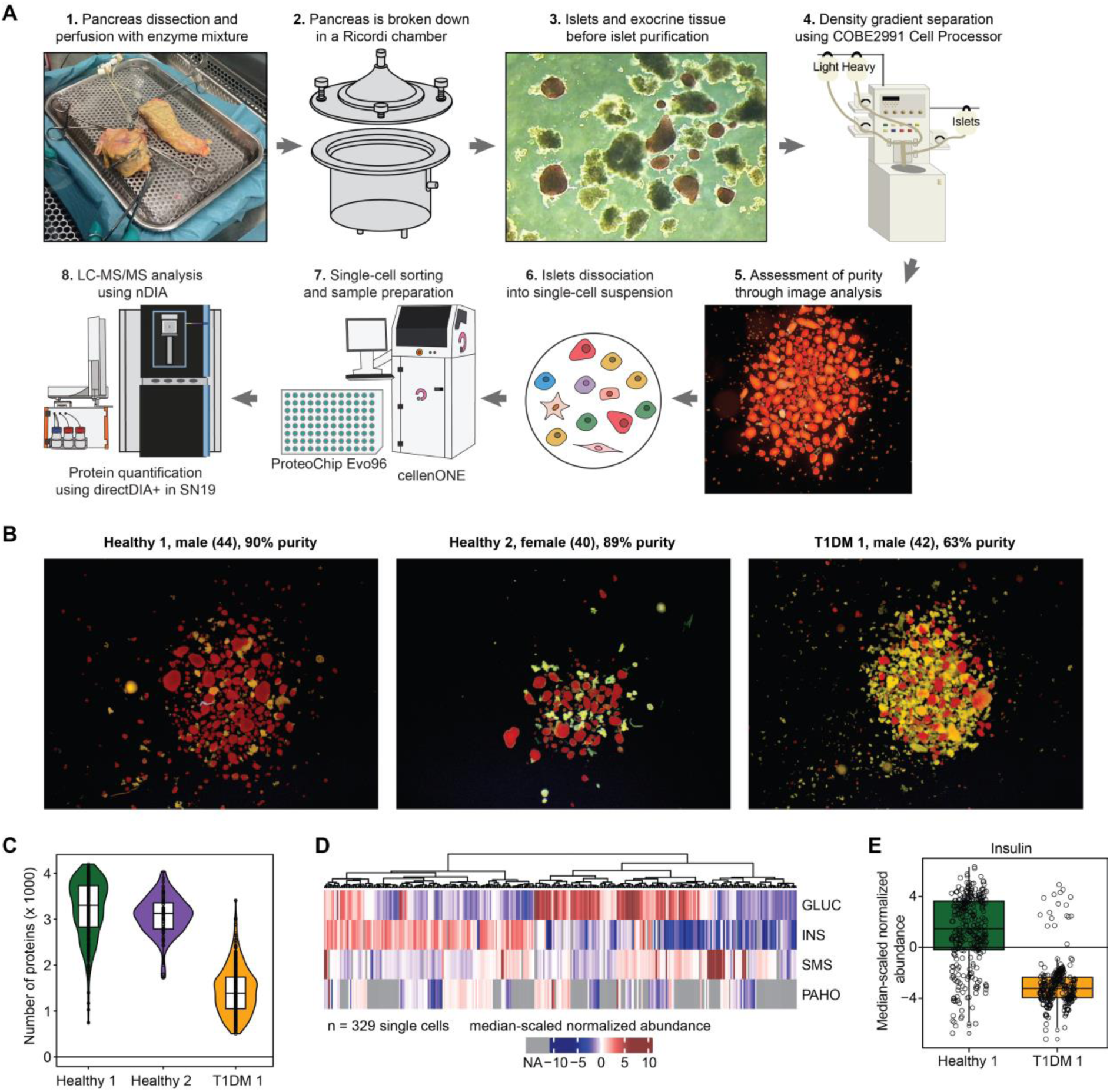
Purification of pancreatic islets and single-cell proteomics analysis. (**A**) Workflow of the islet purification and subsequent single-cell proteomics sample preparation and LC-MS/MS analysis (modified from Shapiro et al.^25^). (**B**) Staining of the purified islets from the corresponding donors, (44 years old), healthy 2 female (40 years old), and T1DM 1 male (42 years old), using dithizone staining (red) highlighting endocrine cells. Exocrine cells remain unstained and appear in yellow. (**C**) Number of proteins quantified per single cell for each donor. (**D**) Heatmap displaying the median-scaled normalized abundance of key pancreatic hormones in single-cell analysis of healthy donor 1. (**E**) Median-scaled normalized abundance of insulin in single cells from healthy 1 and T1DM 1 donors.

To verify the presence of the various pancreatic islet endocrine cell types within our single-cell analysis, we plotted the median-scaled MS-based protein abundance of the main pancreatic peptide hormones glucagon (GCG), insulin (INS), somatostatin (SST) and pancreatic polypeptide (PPY) across the single cells analyzed from healthy 1 (Figure 1D). These hormones are specific to alpha, beta, delta, and gamma cells, respectively. The epsilon cell marker Ghrelin (GHRL) was also detected but at very low abundance and in very few cells. We therefore did not study GHRL in more detail.

The abundance-based hierarchical clustering heatmap shows high, cell-specific expression of each hormone (i.e., elevated levels of only one hormone per cell), with only a few low-frequency exceptions (Figure 1D). The hormone distribution aligns with the expected cell-type composition of a healthy pancreatic islet, with ∼60% beta cells, ∼30% alpha cells, and much smaller proportions of delta (<10%) and gamma (<5%) cells^25^. Surprisingly, we detected insulin in a subset of cells from the T1DM donor at levels comparable to those of a healthy donor, suggesting the persistence of some functional beta cells in these islets (Figure 1E).

Since no established and robustly tested SCP tool is currently available to perform an unbiased cell type annotation, we instead tested established scRNAseq tools for this purpose. While high expression of specific hormones greatly aids in annotating endocrine cells, hormone expression alone is insufficient, as our dataset also includes exocrine cells and potentially rarer cell types such as endothelial and immune cells.

The first SCP analysis of human pancreatic islets was performed by Fulcher et al.^26^ combining mRNA analysis of the same cell to obtain 2-dimensional measurements. The authors used the Seurat-based Azimuth pipeline^27,28^ to integrate their data and successfully annotate cell types using the mRNA measurements first and then protein measurements second to further refine their annotation, demonstrating that protein analysis helps annotating cells by providing complementary information. Here we investigated whether a similar approach could be applied relying on protein abundance measurement only. We analyzed the healthy 1 dataset using the Azimuth app and visualized the output in a Uniform Manifold Approximation and Projection (UMAP) plot (Figure 2A). The Azimuth app identified and classified several cell types of the islet but did not identify delta cells, despite the apparent high level of somatostatin in specific cells from healthy 1 (Figure 1D). A high number of activated stellate and Schwann cells was also assigned which seemed unlikely to be present in such proportions if even present at all.

**Figure 2.**
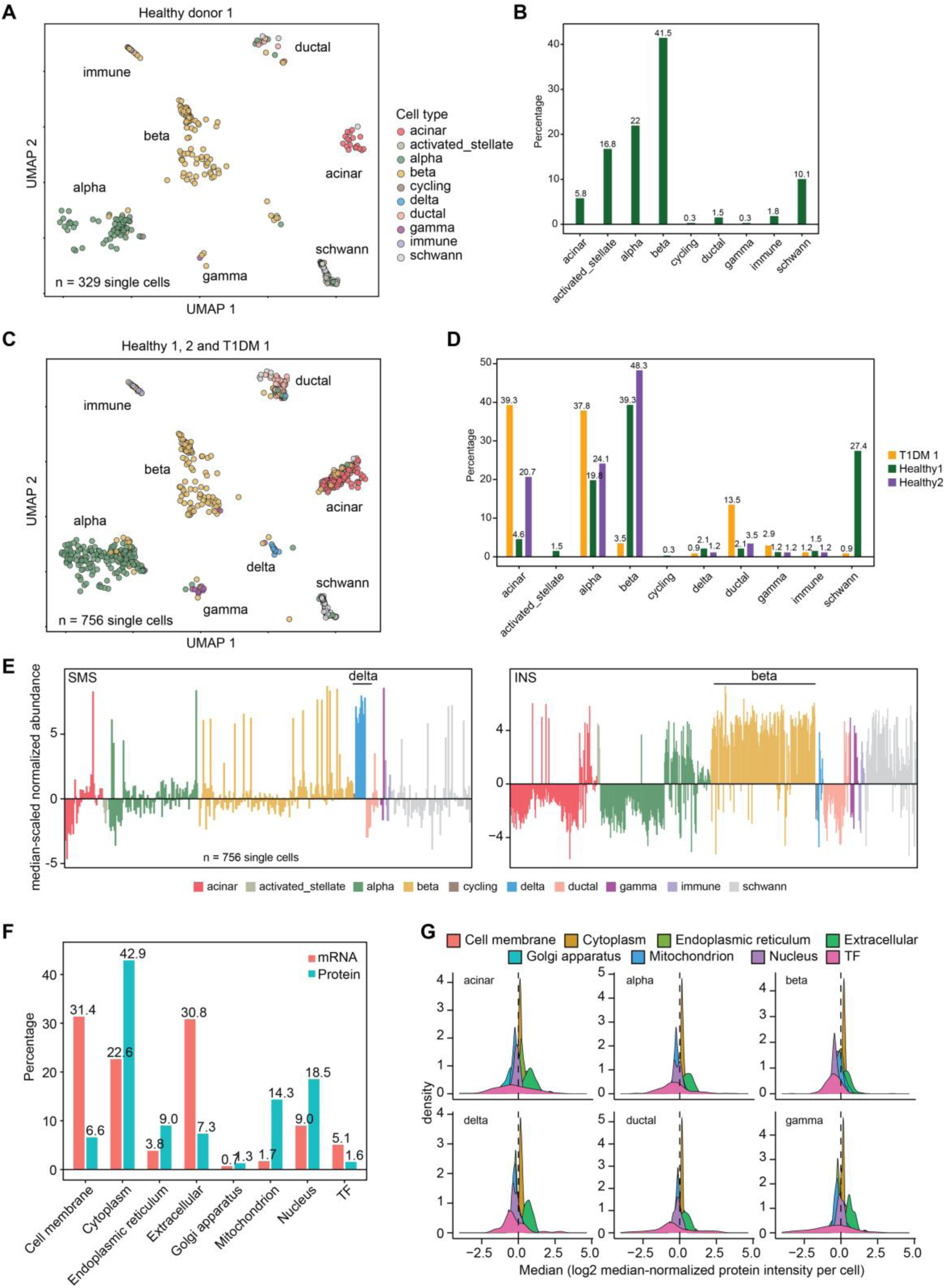
Cell type annotation using scRNAseq tools. (**A**) Uniform Manifold Approximation and Projection (UMAP) projection of the output from the Seurat integration of the Azimuth pipeline of protein abundances of cells from healthy 1 donor. (**B**) Proportion of each cell type annotated via Azimuth in healthy 1 donor. (**C**) UMAP projection of the output from Azimuth after integration of the protein expression values of cells from healthy 1, healthy 2 and T1DM 1 donors. (**D**) Proportion of each cell type annotated via Azimuth in healthy 1, healthy 2 and T1DM 1 donors. (**E**) Median-scaled normalized abundances of somatostatin (SST) (left), and of insulin (INS) (right) across every single cell from healthy 1, healthy 2 and T1DM 1 donors according to each cell type annotated from Azimuth. (**F**) Subcellular annotation of proteins from our dataset and protein from the corresponding mRNA obtained from the SingleR reference pancreas dataset, using Deeploc 2.0 for subcellular annotations and gene ontology (GO) annotation for proteins involved in “DNA-binding transcription factor activity” (TF).

We then tested whether Azimuth could integrate data from the three datasets combined. This could prove difficult as the datasets were acquired at different dates and present differences in analytical depth, particularly the T1DM 1 dataset. However, the cell proportions in each sample appear to partially reflect the expected composition of pancreatic islets (Figures 2C and 2D). Notably, T1DM 1 cells only contained a few cells identified as beta cells (3.5%) and a high proportion of acinar and ductal cells with 39.3 and 13.5%, respectively. The proportion of exocrine cells was roughly in line with the expected purity of the T1DM 1 islet extraction (Figure 1B). Schwann cells were again overrepresented in the Healthy 1 sample, accounting for 27.4%, which seems unlikely given that these cells are typically located at the islet periphery, while the endocrine tissue proportion was ∼90% in this extraction.

Azimuth relies on so-called anchor genes to help define cell populations subsets, with each cell type represented by 10 gene anchors^28^. These anchors were defined by integrating multiple reference scRNAseq datasets from various sources. Therefore, we studied the expression of these markers in our dataset. Many of the proteins corresponding to these gene anchors were not found in our dataset (Supplementary Figure 1). Particularly, we detected none of the corresponding proteins for Schwann cells, which is likely because we had too few or no Schwann cells at all in the datasets. This is surprising since a high proportion of these cells was detected in healthy 1 donor and is likely an issue from attempting to annotate SCP data with a tool meant for scRNAseq analysis. This was even more apparent when plotting the abundance of pancreatic hormones since most cells annotated as Schwann cells exhibit high levels of insulin and are likely beta cells (Figure 2E). In addition, many cells harboring high abundance of somatostatin were not annotated as delta cells.

This prompted us to analyze method sampling bias in scRNAseq and SCP. The mRNA approaches can be biased toward certain mRNA lengths but can detect low abundant transcription factors and cell surface marker proteins. Conversely, transmembrane-spanning as well as chromatin-interacting and very low abundant protein remain challenging in proteomics due to suboptimal extraction and digestion for the former and to the lack of amplification for the latter. This bias analysis showed that the scRNA-seq dataset we examined contained a higher proportion of transcription factors and membrane proteins, as determined using the reference pancreatic islet dataset from SingleR (Figure 2F).

While our SCP analysis showed a higher proportion of cytoplasmic proteins. An analysis of the distribution of single-cell protein abundances using DeepLoc, a deep learning tool to predict subcellular localization of eukaryotic proteins, showed that proteins from the extracellular compartment and cytoplasm are the most abundant while transcription factors present the widest distribution of protein abundances, including some of the lowest expressed proteins (Figure 2G). Since transcription factors and cell surface membrane-spanning proteins are often used as reference markers in scRNAseq studies and in flow cytometry, our observations stress the need to extend the number of reference markers for a given cell type, including more cytoplasmic proteins as well as to define SCP-specific markers. This would also greatly facilitate future multi-OMICS single-cell analysis studies to corroborate and integrate results from multiple OMICS sources.

In addition to analytical bias, some of the markers annotated as specific for a cell type were not specific at the protein level in our datasets. For instance, PCSK, the gene that encodes for the enzyme prohormone-convertase 1, is often used as a marker of delta cells including in Azimuth but is also involved in proinsulin processing and present at similar levels in beta cells (Supplementary Figure 2). The correlation of PCSK1 is also higher with insulin (0.37) than with somatostatin (0.12). This discrepancy was particularly apparent in less abundant cell types, such as delta and gamma cells, where we could only confirm a few markers.

PCSK1 was not an isolated case, and it appears that many markers both differed by their absence, but also by their specificity at the protein level. For epsilon cells markers we detected ghrelin in very few cells as the sole marker; only one marker of endothelial was detected and widely expressed; 3 markers were measured for immune cells and were not in agreement with each other; 3 markers for quiescent stellate cells with two of them being relatively widely expressed among other cells, which was also the case for activated stellate with three markers; two in gamma cells including tubulin beta-2A (TUBB2A), which was mostly expressed in beta cells. While five markers were detected in delta cells, PCSK1 was as high in beta cells as mentioned above, which was the same for alpha-1-antitrypsin (SERPINA1) and signal peptidase complex catalytic subunit SEC11C. Only retinol-binding protein 4 (RBP4) matched cells with high levels of somatostatin. Four markers were detected in alpha cells, with transthyretin (TTR) and prohormone-convertase 2 (PCSK2) showing good agreement with glucagon (GLUC) expression levels, which was not the case for protein phosphatase 1 regulatory subunit 1A (PPP1R1A). For beta cells, the detected markers seemed in agreement with insulin abundance including the levels for hydroxyacyl-coenzyme A dehydrogenase (HADH), gelsolin (GSN), glucose-6-phosphatase 2 (G6PC2), neuronal pentraxin-2 (NPTX2) and islet amyloid polypeptide (IAPP). Ductal cells were mostly characterized by the expression of annexin A4 (ANXA4), and other detected markers were expressed in other cell types. Acinar cells had three markers present at high abundance in other cell types including carboxypeptidase A1 (CPA1), chymotrypsinogen B2 (CTRB2) and chymotrypsin-like elastase 3A (CELA3A). Overall, this highlights the discrepancies between mRNA and proteins levels since mRNA markers specific for certain cell types do not necessarily translate to the same specificity at the protein level.

This also demonstrates that while tools such as Azimuth can be applied to SCP-only data, they should be used cautiously and complemented with other approaches for cell type annotation, as scRNA-seq and SCP data differ in both quality and quantity (i.e. dynamic range). In the study from Fulcher et al.^26^, it was optimally used though as the mRNA readings were employed for cell type annotation.

To overcome this issue, we curated a reduced list of markers from Azimuth and the literature, as well as some that were manually confirmed through correlation analysis with pancreatic hormones (Figures 3A). We exploited this marker list in two ways, first by determining the specificity of each cell toward the marker list of the corresponding cell type. Secondly, we also performed a marker-based similarity approximation using SingleR by creating a dummy with binary expression based on the marker list. Finally, we combined these two annotations with Azimuth annotations to provide the final cell type assignment. Endothelial, Schwann and epsilon cells were excluded from the analysis as we could not reliably quantify markers from these cells, suggesting that they were either too few or absent from the dataset. Similarly, we could not differentiate between activated or quiescent stellate cells, so these two categories were grouped under the “stellate” term, which only accounted for a handful of cells in the entire dataset. The resulting annotations were in better agreement with known markers and particularly with peptide hormones. The average marker abundance showed high specificity within each cell type (Figure 3A right panel). Delta cells were also annotated in the healthy 1 dataset contrary to when using Azimuth only.

**Figure 3.**
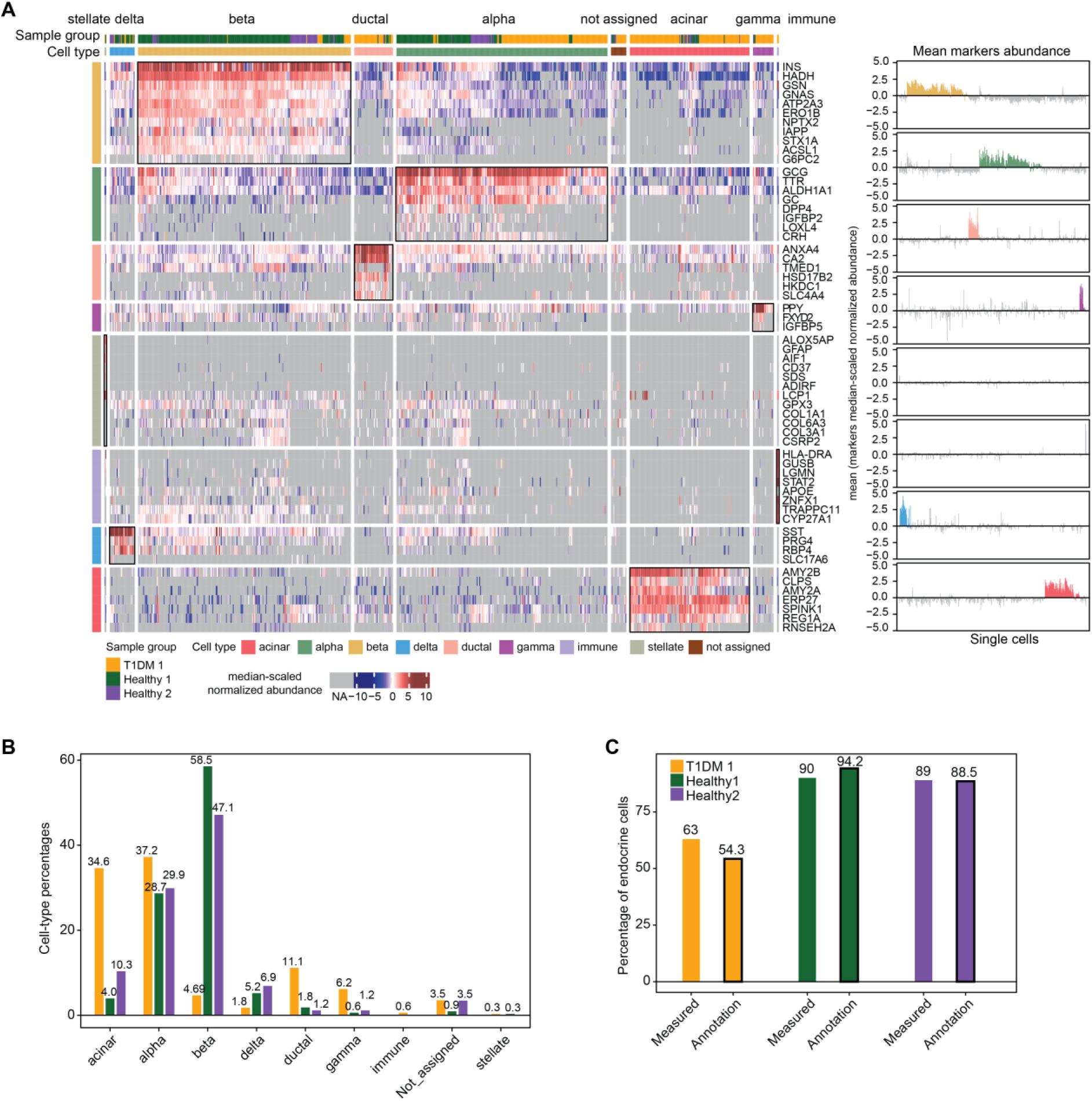
Cell type-specific markers and combined cell type annotations. (**A**) Heatmap of median-scaled normalized abundance of the curated markers across all cells in healthy 1, healthy 2 and T1DM 1 donors, classified by cell type following our annotation strategy (left). (right) Distribution of the average abundance for each marker group within the corresponding cell type across every cell. (**B**) Proportion of each cell type in healthy 1, healthy 2 and T1DM 1 donors. (**C**) Percentage of endocrine cells measured with staining and microscopy analysis from Figure 1A compared to the percentage from our annotation (black contour).

Importantly, the number of proteins quantified per cell did not appear to bias cell type annotation neither with Azimuth nor with our approach as exemplified in the T1DM 1 dataset. Therefore, deep proteome coverage is not crucial for annotating cells from pancreatic islets. The estimated proportions matched better with expected cell proportions in healthy donor 1 islets (Figure 3B). Lastly, the purity (% of endocrine cells) was also in line with microscopy-based measurements (Figures 1B and 3C), showing that our single-cell preparation and sorting did not result in a cell type bias among endocrine and exocrine cells. Finally, the presence of beta cells in T1DM 1 islets was further confirmed by our combined annotation and by the presence of key beta cell markers in addition to insulin.

This newly annotated dataset allowed us to compare findings with Fulcher et al.^26^ (Supplementary Figure 2). We hereby confirmed that aldehyde dehydrogenase 1A1 (ALDH1A1) and gelsolin (GSN) showed higher levels in alpha cells, and hydroxyacyl-coenzyme A dehydrogenase (HADH), ERO1-like protein beta (ERO1B), syntaxin-1A (STX1A) and neuroendocrine secretory protein 55 (GNAS) were higher in beta cells compared to other cell types. However, the abundances of endoplasmic reticulum resident protein 29 (ERP29) and endosome/lysosome-associated apoptosis and autophagy regulator 1 (ELAPOR1) were not strikingly higher in alpha cells than in beta cells and while the calmodulin regulator protein PCP4 (PCP4) showed high abundance in delta cells, it was also highly abundant in beta cells.

Interestingly, some of the cells annotated as beta cells show low levels of insulin compared to other beta cells, but similar levels of other beta cell markers. These cells likely correspond to highly degranulated beta cells which have secreted their insulin content. Identifying these cells through the abundance of other markers highlights the strength of the analysis as well as the potential of SCP to detect such differences, which would not be visible at the mRNA level. We also detect multi-hormone cells i.e. annotated beta cells with relatively high levels of glucagon and conversely alpha cells expressing insulin. Although some of these cells may have been processed with unseen debris of another cell type present in the cell suspension, multi-hormone cells have previously been reported even in recent studies^29^. Some gamma cells also harbored insulin or glucagon, which has been reported previously^30^.

Obtaining pre-annotated dataset is key to abundance-based annotation transfer. To test the usefulness of our approach and of our dataset we annotated another dataset from a healthy donor (healthy 3, female) using the combined Azimuth and marker-based annotations. We then compared it to a machine learning (ML) approach where we trained a random forest model on our combined annotated dataset (healthy 1, healthy 2 and T1DM 1) and applied the trained model to annotate the new dataset (healthy 3). We considered the combined annotation that we previously established as the ground truth for the sake of comparison, bearing in mind that some cells could not be annotated (NA) or could have been misidentified. Although this dataset was of lower proteome depth compared to the other two healthy datasets (Supplementary Table 1), it did not hamper cell type annotation (Figure 4A). The ML model displayed 91% accuracy compared to our combined annotation, when using the three datasets including annotation of the NA, which otherwise would have only resulted in five cell transition to another cell type out of 75 cells.

**Figure 4.**
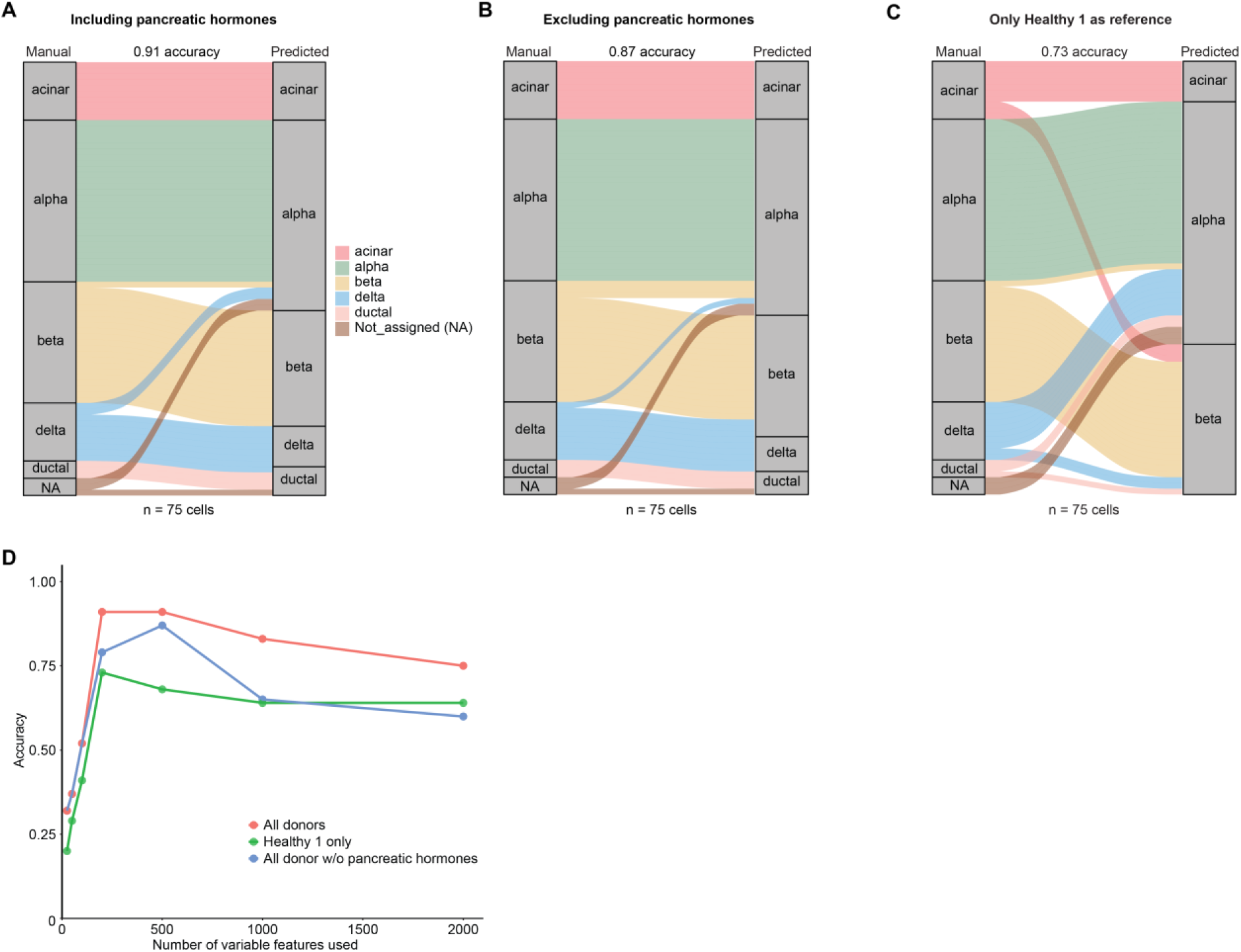
Machine learning approach to cell type annotation based on our reference dataset. (**A**), (**B**), and (**C**) alluvial plots representing the initial cell type annotation from our previous approach (manual, left on the plots) and after applying random forest models based on the following SCRAN-normalized datasets: the whole dataset (500 features), the whole dataset but excluding pancreatic hormones (500 features), healthy 1 donor dataset only (200 features). The accuracy of the annotation was also calculated compared to the “ground-truth” annotations. The number of features selected to train the model was based on the best results in terms of accuracy. (**D**) Accuracy of the annotation depending on the number of variable features from the reference dataset selected to train the model.

Pancreatic hormones are known to be highly expressed and in some cases their mRNA have been shown to represent up to 50% of the mRNA pool. Therefore, we calculated the average rank and abundance of each protein markers used to annotate the cells including each pancreatic hormone in the corresponding cell type (Supplementary Figure 3). Even though pancreatic hormones did not represent 50% of the total protein intensities, they generally ranked among the top 10 most abundant proteins in the corresponding cell types, questioning the impact of these proteins in annotating cell types. Consequently, to test their impact on cell type annotation, we removed each pancreatic hormone from our reference dataset before training and from healthy 3 test dataset prior to applying the trained model. The model was close in accuracy to the model incorporating pancreatic hormones (91% vs 87%), suggesting that even in absence of these highly expressed and cell-type specific proteins, the proteome signature from three different datasets despite batch effects still enables high quality annotation transfer using a ML approach (Figure 4B). Finally, we trained the ML model on healthy 1 dataset only. The accuracy highly dropped from 91 to 73%, emphasizing that the size of the training dataset is important for establishing the most accurate models and that only one of our datasets is insufficient (Figure 4C).

We also evaluated how the number of variable features affected ML model performance by selecting the top 25 to 2000 proteins with the highest variance in the combined dataset and comparing annotation accuracy, including conditions where pancreatic hormones were excluded or only the Healthy 1 sample was used for training (Figure 4D). Accuracy peaked at ∼500 features for the combined datasets and after removing pancreatic hormones, whereas for the Healthy 1 dataset alone it peaked at ∼200 features (also shown on Figures 4A-C). The curve also stayed much higher when using the three combined datasets for the training of the model. Importantly, the accuracy was much lower at 1000 and 2000 features. This demonstrates that a limited number of features is sufficient to confidently annotate cell types from a reference proteome dataset. This is also in line with the rank of cell type-specific markers, which ranked high in general and mostly in the top 500-1000 proteins (Supplementary Figure 3). This likely explains why cell-type annotation is not strongly influenced by proteome depth, as most cells we analyzed had over 1,000 proteins quantified. It also demonstrates that our dataset can serve as a reference for annotating future pancreatic islet datasets.

After exploring cell type annotations and confirming their robustness, we compared the various endocrine cell types by combining the healthy 1, healthy 2 and T1DM 1 donor datasets, and applying NEgative Binomial mixed model Using a Large-sample Approximation (NEBULA)^31^. NEBULA accounts for sampling depth as well as batch effect by integrating it in the differential analysis. This proved particularly useful in our datasets as some cells are much lower represented such as delta and gamma cells and combining different batches helps improve the analysis. This allowed us to compensate for the difference between donors as well as technical variations to highlight specific differences between the various cell types (Figure 5A). The estimated log2 fold-changes (FCs) and p-values highlighted key proteins, which could be included in our protein marker list showing promising specificity. Those include clusterin (CLU), sulfide quinone oxidoreductase (SQOR), aromatic-L-amino-acid decarboxylase (DDC), and dipeptidyl peptidase 4 (DPP4) for alpha cells; palmitoyl-protein thioesterase ABHD10 (ABHD10), ankyrin repeat and SOCS box protein 9 (ASB9), DNA dC->dU-editing enzyme APOBEC-2 (APOBEC2), Synaptic vesicular amine transporter (SLC18A2), carboxypeptidase E (CPE), ganglioside-induced differentiation-associated protein 1 (GDAP1), retinol dehydrogenase 14 (RDH14), cell adhesion molecule 1 (CADM1), complexin-2 (CPLX2), cytoplasmic glycerol-3-phosphate dehydrogenase (GPD1), alpha-1,3/1,6-mannosyltransferase ALG2 (ALG2), and golgi-resident adenosine 3’,5’-bisphosphate 3’-phosphatase (BPNT2) for beta cells; embryonal fyn-associated substrate (EFS) for delta cells; nesprin-2 (SYNE2), insulin-like growth factor-binding protein 7 (IGFBP7) and glutathione peroxidase 1 (GPX1) for gamma cells (Supplementary Figure 4).

**Figure 5.**
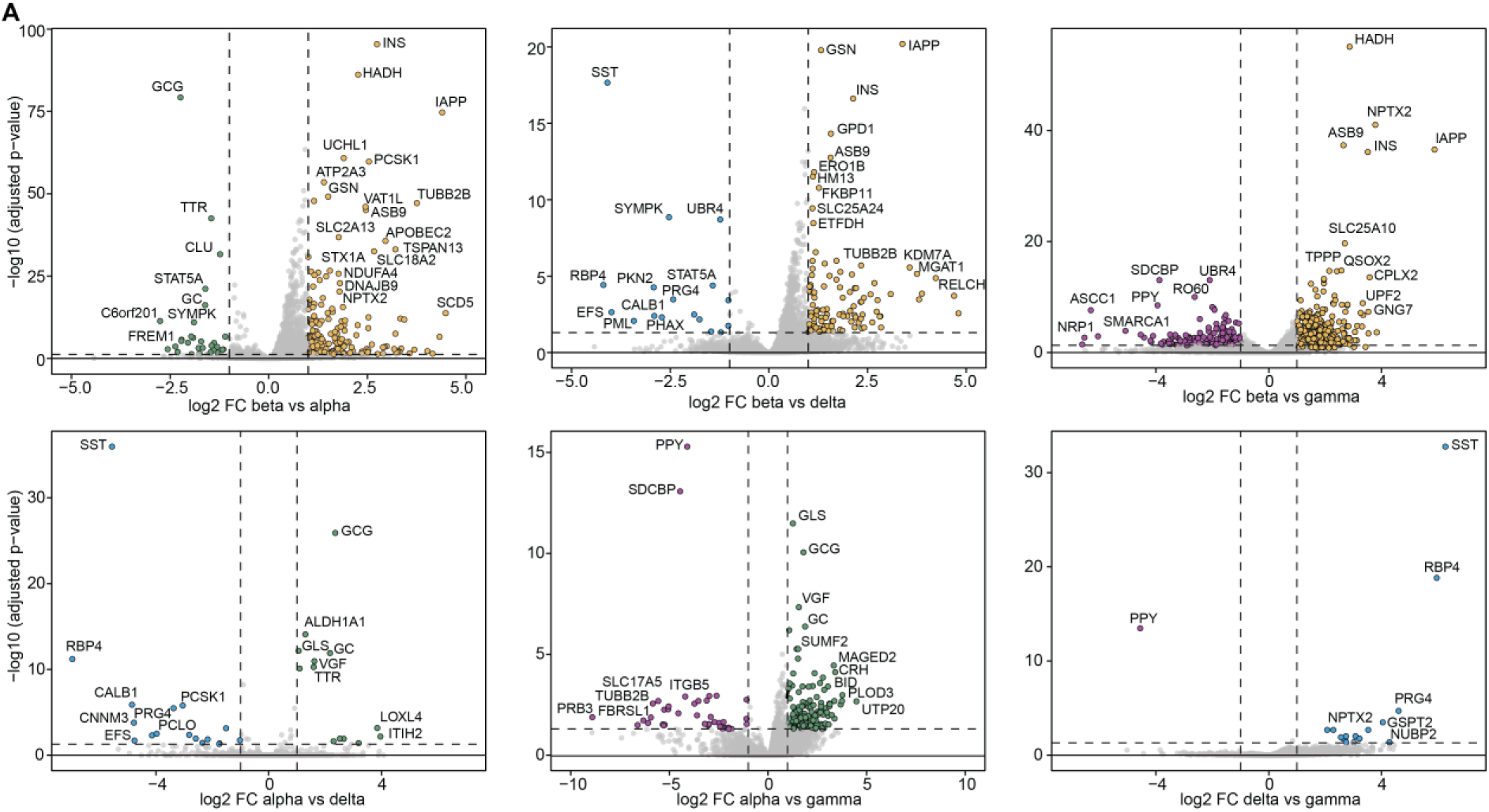
Endocrine cell-type comparisons. Volcano plots presenting the paired differential expression analyses using NEBULA between alpha, beta, delta and gamma cells, using the combined data from healthy 1, healthy 2 and T1DM 1 donors. p-values were corrected using Benjamini-Hochberg method. Highlighted proteins showed abs (log2 FC)>1 and Benjamini-Hochberg adjusted p-value <0.05. Protein with higher abundance in one cell type over the other are highlighted in green, yellow, blue, and purple for alpha, beta, delta and gamma cells, respectively.

Strikingly, beta cell proteomes showed significantly more differential proteins than the other cells. In part this is most likely due to the difference in cell numbers (higher number of beta cells), However, this is not the only explanation, since the numerical differences between alpha and delta or gamma cells are also large, yet the number of significantly regulated proteins is much lower than in beta cells (Supplementary Table 1). Notably the number of proteins expressed in beta cells was higher on average than in the other cell types (Supplementary Figure 5A). As a result, only proteins significantly upregulated in beta cells showed a GO enrichment for pathways related to regulation of hormone levels and insulin secretion (Supplementary Figure 5B). This is likely because beta cells are extremely active within the islets. Surprisingly, beta cells also had the highest number of proteins expressed on average in the T1DM 1 donor (Supplementary Figure 5A). This should be taken with caution though as the number of beta cells in the T1DM 1 dataset was in general very low (n=16).

Finally, having established and identified cell types in our datasets, we compared population subsets in healthy 1 and T1DM 1 donors to highlight specific features of diabetes (Figure 6A). Since, the analysis of the different donors was done at a different time, creating technical variability, this effect could not be fully removed and is in part confounded with the biological variation i.e. T1DM or donor to donor differences. Therefore, we employed two different strategies to compare T1DM 1 against healthy donors, and healthy 1 against healthy 2. The first strategy was to use alpha cells as a reference in each dataset to compare their differences with beta cells between donors. While this approach would not enable us to detect effects that evenly affect cells within the pancreatic islet, this would still allow us to pinpoint specific differences to beta or alpha cells. To avoid large differences due to analytical depth, we only considered proteins which were present in at least 50% of each cell type in both donors of each paired comparison. Surprisingly, the correlation was relatively high (*r* = 0.7) considering that the samples were not only from different batches but also from healthy and T1DM donors. As expected, the shared upregulated proteins in beta cells include some of our selected markers such as IAPP, NPTX2, HADH and insulin (INS) as well as other proteins involved in insulin metabolism such as PCSK1, while downregulated proteins corresponded to markers of alpha cells such as GCG, ALDH1A1, TTR and GLS. The proteins that were uniquely significant in T1DM comparison includes neudesin (NENF), which is downregulated and is known to play a role in insulin resistance in T2D. The HIG1 hypoxia inducible domain family member 1A (HIGD1A) was much lower in T1DM and is protecting cells from hypoxia. Pterin-4-alpha-carbinolamine dehydratase (PCBD1) was lower in beta cells and is known to cause early onset of a form of non-autoimmune diabetes. Many more proteins were uniquely significant in healthy 1. However, when investigating their FCs, most also showed a similar FC change in T1DM but did not pass the p-value threshold likely due to the very reduced size of the beta cell population in the T1DM group.

**Figure 6.**
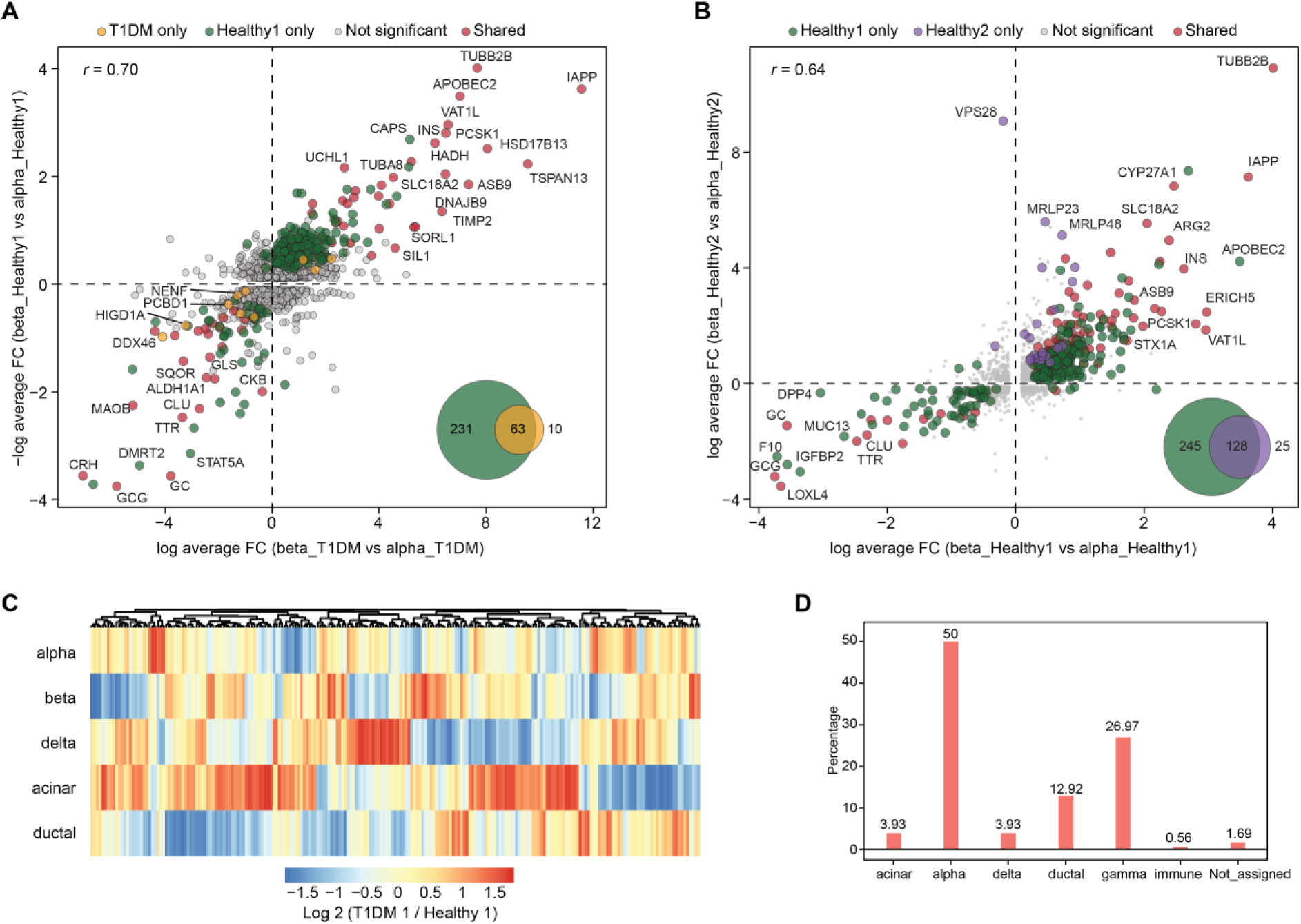
Single-cell proteomics provides insights into type 1 diabetes and donor-specific cell-type differences. (**A**) Differential analysis of beta cells versus alpha cells using Wilcoxon rank sum test on Seurat SCTransform-normalized and integrated data from healthy 1 (y-axis) and T1DM 1 (x-axis) donors. FC are plotted as log average of beta – alpha. P-values were adjusted using Bonferroni correction and proteins showing abs(ln FC)>1 and adjusted p-value<0.05 are highlighted. Correlations were calculated using Pearson’s method. (**C**) Heatmap of log2 FC of significant proteins between healthy 1 and T1DM 1 donors in alpha, beta, delta, acinar and ductal cells. FCs and p-values were obtained from differential expression analysis using edgeR. Counts were normalized to account for sequencing depth and composition bias (TMM normalization). Batch effects were accounted for by including batch as a covariate in the generalized linear model. Differential expression was assessed using a likelihood ratio test. FC and p-values were calculated using a generalized linear model likelihood ratio test. Proteins expression was filtered for proteins present in at least 50% of the cells of each batch in a given cell-type. To compensate for batch effect, significant proteins were further filtered and only proteins showing abs (log2 FC)>1 and adjusted p-value<0.05 in one cell type only were considered specific and presented in this heatmap.

Strikingly, we did not record more differences between healthy 1 and T1DM 1 donors than between healthy 1 and healthy 2 donors suggesting that the effect of T1DM is either not chronic on the remaining functioning beta cells or that it is systemic in the islet (Figures 6A and 6B). The donors were in a similar age group so this factor should not have influenced these observations (42, 44, 40, for T1DM, healthy 1, healthy 2), but healthy 2 was female, which could impact the comparison. However, the changes in healthy 2 had lower FC differences with healthy 1 compared to T1DM 1 and affected mostly mitochondrial proteins involved in cellular respiration and upregulated in beta cells, suggesting a slight difference in mitochondrial function and metabolism between beta and alpha cells in healthy 2 compared to healthy 1 (Supplementary Figure 6). The relatively good correlation of the FCs show that the data of all three donors are in agreement for major markers but still retain levels of disease and donor-associated differences.

The absence of large outliers may also underline that the disease was well established in the T1DM 1 donor, and the peak of the inflammation and immune reaction is already largely gone leaving the remaining beta cells with their normal function but in greatly reduced numbers.

To further investigate differences between T1DM 1 and healthy donors, we applied a second approach where the data was analyzed with edgeR instead of NEBULA as the number of cells in some cell types was too low. In this framework, batch effects were accounted for by including batch as a covariate in the generalized linear model, rather than applying an explicit batch-correction algorithm. This approach preserves biological variation while adjusting for analytical depth and batch-specific biases. We selected the most abundance cell types, namely alpha, beta, delta acinar and ductal and performed paired comparisons for each cell type between T1DM 1 and healthy 1 donors, only considering proteins which were present in at least 50% of cells within a cell type in each donor. Then, to further remove proteins which were systematically regulated in multiple cell type, only proteins which were significantly different in one cell type only were considered as specific feature of this particular cell type.

Finally, we analyzed pancreatic islets from a second donor (T1DM 2). Unfortunately, no cells were annotated as beta cells among the 392 cells which we analyzed (Figure 6D), which was reinforced by the absence of detection of insulin (Supplementary Figure 7). However, the number of gamma cells identified was surprisingly high (∼25% of the total cell counts). The proportion of gamma cells in T1DM 1 donor was also high in comparison to healthy donors (∼6%), and while much lower than in T1DM 2, this is also because ∼45% of the cells were from exocrine tissue so the actual value would be >10% among islet cells only. This large difference in proportion of gamma cells was not observed for other cell types and only alpha cells showed a slight increase in proportion in diabetic islets. This observation has not been made before and may be a consequence of diabetes.

## Discussion

Our study represents an early attempt to obtain single-cell pancreatic islet proteomes from multiple fresh (i.e. non-frozen) cells from donors’ organs, representing a considerable technical challenge. Unpredictable donor availability, combined with challenges in cell storage for single-cell proteomics, prevented us from performing SCP analyses simultaneously, which would have reduced technical variability that partly overlaps with biological differences in our dataset. Although strategies such as those applied in this study can partially mitigate this issue, SCP would benefit greatly from the development of optimized sample storage methods. To date, most efforts have focused on freezing and fixation^32–35^, where both demonstrated limitations with risk of altering the cells and proteome during long-term storage.

We readily identified alpha, beta and delta cells based on their specific proteome profiles as these were the most abundant cell types in the analyzed pancreatic islets. Known markers such as glucagon for alpha cells, insulin for beta cells and somatostatin for delta cells were among the most abundant proteins detected in the respective cells. Moreover, we identified additional sets of protein markers, which were unique to these three cell types. We also highlighted few additional markers for the rarer cell types such as gamma cells. These cells would greatly benefit from a reanalysis using cell enrichment strategies to artificially increase their numbers. While we detect ghrelin, the lack of epsilon-specific markers failed to clearly identify epsilon cells. This will require larger datasets of high quality, which needs to be considered while planning future studies. Nevertheless, we demonstrated that combining datasets with different biological meaning is possible and enabled us to annotate cell types from these datasets as well as perform differential expression analysis. Adding more dataset to this work will contribute to increasing its relevance.

Since only one other SCP study of pancreatic islets is currently available^26^, and it included fewer cells and lower analytical depth, we relied on scRNA-seq tools. However, generating such SCP datasets will be crucial, as we show that cell type annotation differs substantially and that many markers identified as specific in scRNA-seq studies are either undetected due to analytical bias or insufficiently specific at the protein level.

While extremely challenging, collecting SCP data of T1DM pancreas at various stages of the disease would greatly help recapitulate the disease progression and further provide clues on its development. Our study demonstrates that this limitation is no longer a technical issue in SCP but rather a consequence of the limited availability of such samples.

Nevertheless, this study and data can serve as a reference for future single-cell proteomics studies of pancreatic islets and T1DM.

## Acknowledgements

Work at The Novo Nordisk Foundation Center for Protein Research (CPR) is funded in part by a donation from the Novo Nordisk Foundation (NNF14CC0001, NNF24SA0098829 and NNF21OC0072070). This project was supported by a center-of-excellence grant from the Danish National Research Foundation to Copenhagen Center for Glycocalyx Research (DNRF196). This project was also supported by a grant from the Danish Agency of Higher Education and Science to establish the PLATO research infrastructure: Danish National Mass Spectrometry Platform for Proteomics and Biomolecular Imaging (5229-00012B). J.V.O. and M.L. are also funded by Novo Nordisk a/s (CELFFI-2022-002843). P.S. is funded by the Swedish Research Council (2022-00323) and by the Åke Wiberg foundation (M24-0238). The study was supported by the Swedish Research Council (VR 2022-01185, VR 2023-02221) the Ernfors Family Fund, Nils Eric Holmstens forskningsstiftelse, Barndiabetesfonden, Diabetesfonden, the governmental strategic research organization-SFO StemTherapy (K.-H.G.), and ExoDiab.

## Author contributions

P.S., L.G., O.K. and J.V.O. designed the experiments. P.S., M.L. and L.G. prepared all samples and performed proteomics experiments. P.S., L.G. and M.L. analyzed the resulting data. P.S., L.G., O.K. and J.V.O. wrote the first draft of the manuscript. S.R. and K.-H.G. critically evaluated the results. All authors read, edited and approved the final version of the manuscript.

## Material and Methods

### Ethics

Pancreases from heart-beating organ donors treated as intended for organ transplantation were procured through the Nordic Network for Clinical Islet Transplantation (https://nordicislets.medscinet.com/en.aspx). Consent for organ donation (for clinical transplantation and for use in research) was obtained via online database (https://www.socialstyrelsen.se/en/apply-and-register/join-the-swedish-nationaldonor-register/) or verbally from the deceased’s next of kin by the attending physician and documented in the medical records of the deceased in accordance with Swedish law and as approved by the Swedish Ethical Review Authority (Dnr 2023-01845-01). All tissue included in the study was procured, stored and analyzed as approved by the Regional Ethics Committee in Uppsala (Dnr: 2015/444).

### Human Pancreatic Islet Isolation

Pancreases from 3 brain-dead organ donors were used for islet isolation (donor characteristics displayed in Supplementary Table 2). The islets were isolated according to clinical routine as previously described^36^ at Rudbeck Laboratory, Uppsala Sweden. In short, pancreases were perfused intraductally using a mixture of collagenase and thermolysin (Supplementary Table 3). The organ was digested and the islets separated from the exocrine tissue using a continuous gradient purification using a COBE 2991 Cell Processor (Cardian, Lakewood, CO, USA). Islet yield, expressed as islet equivalents (IEQ), and purity were determined in a standardised procedure using a digital analysis system (Cellimage, Uppsala, Sweden) that discriminates dithizone-positive islets from unstained exocrine particles^37^. Isolated islets were kept under optimal culture conditions; in semi-permeable bags in 25°C incubator, with media changes every second day, until dissociation to single cells. Three tissue samples were also taken and preserved in either N^2^ or formalin from all donors.

Quality tests were performed on the islets the day after islet isolation using a dynamic perifusion system (Biorep Technologies, TP software Version 4.7, cRIO software Version 4.7. Miami Lakes, Florida, USA). Duplicate aliquots of 20 hand-picked islets were placed in filter-closed chambers and perifused sequentially with a buffer containing low (1.67 mM), high (20 mM) and then low (1.67 mM) glucose. Following perifusion, insulin ELISA was performed according to the manufacturer’s instructions (Cat. No. XXX, Mercodia, Uppsala, Sweden) to determine the insulin levels in the effluent. The glucose-stimulated insulin response was expressed as a stimulation index (SI), which was defined as the ratio between the areas under the curves that were calculated for the low and high glucose concentrations. The results yielded high SI for both healthy donors, and unsurprisingly, a very weak response from the donor with T1DM (Supplementary Table 2).

### Single-cell dissociation

From each donated pancreas, at least 3000 IEQ were collected from an endocrine tissue isolate (endocrine content of ∼90% based on automated image analysis after dithizone staining, T1D donor 63 % purity). The islets were dissociated into single cells by adding 1 mL of pre-heated (37°C) Accutase Cell Dissociation Reagent (Gibco, A11105-01) containing 10 mg/mL DNase (Pulmozyme, Gentech), while pipetting up and down for 1-2 min. Non-dissociated islets were allowed to settle in the incubator for 5 min. The single-cell-containing supernatant was removed, and 1 mL of fresh Accutase-DNase mix was added to the remaining islets again while pipetting up and down 3-4 times. The cycle was repeated until no sedimented islets were visible. The single-cell suspension was centrifuged, the Accutase-DNase mix removed, and the cells were washed with sterile DPBS (pH=7.3). The cells were spun down again and filtered through a 40 µm filter. After a final centrifugation, the DPBS was replaced with degassed PBS. Immunofluorescent staining of endocrine cells Paraffin sections (6 µm) from the T1DM tissue sample block were processed using a standard technique. All antigens were unmasked by heat-induced antigen retrieval using citrate buffer according to the manufacturer’s instructions (Invitrogen). The sections were blocked with 5% normal goat serum (Agilent) for 30 min and incubated with anti-insulin, anti-glucagon, and anti-somatostatin antibodies (Supplementary Table 4) diluted in 5% goat serum (in 1× TBS-Tween) for 60 min RT. Nuclei were stained with 500 nM SYTOX Orange 5 min RT. The slides were examined and photographed using OLYMPUS BX43.

### Cell isolation and single-cell sample preparation

Cells were detached using trypLE select (Gibco), fresh medium was added to dilute the tryplE and cells were centrifuged at 400 x g for 3 min then washed twice with Phosphate Buffered Saline (PBS) (Gibco) with centrifugation in-between, before being resuspended in degassed PBS with thorough pipetting. The cell concentration was adjusted to ∼200 cells/μl with degassed PBS. 1 cell, or 20 cells were sorted into Evo96 proteoCHIPs (Cellenion) onto pre-dispensed lysis and digestion buffer consisting of 0.2% n-Dodecyl-β-D-Maltoside (DDM), 100 mM TEAB, 10 ng/μl lysyl endopeptidase and 20 ng/μl trypsin. The digestion was conducted at 50°C for 1.5 hours and quenched by adding 4 μl of 0.1% formic acid to the wells. Lastly samples were loaded onto pre-equilibrated Evotips (Evosep) following manufacturer’s protocol and stored in the fridge prior to LC-MS/MS analysis.

### LC-MS/MS analysis

Evotips containing the samples were loaded on an Evosep One (EvoSep Biosystems) LC system connected to an Orbitrap Astral MS (ThermoFischer Scientific). Samples were analyzed in 40SPD (31-min gradient) using a commercial analytical column (Aurora Elite TS, IonOpticks) interfaced online using an EASY-Spray source. The Orbitrap Astral MS was operated at a full MS resolution of 240,000 with a full scan range of 380 − 980 m/z when stated. The full MS AGC was set to 500% or 30 ms. MS/MS scans were recorded with 4 Th isolation window, 6 ms maximum ion injection time. MS/MS scanning range was from 380-980 m/z were used. The isolated ions were fragmented using HCD with 27% NCE.

### MS data analysis

Protein identification and quantification was performed with Spectronaut v 19 (Biognosys) applying the spectral library-free approach (directDIA +) using the human protein reference database (Uniprot 2022 release, 20,588 sequences) complemented with common contaminants (246 sequences). Default settings were used, except that carbamidomethylation was not included in the fixed modifications and cross-run normalization was also removed. Data were analyzed using R.

### Data analysis

The data was processed using R and R Studio, and the packages Seurat, Nebula, SCRAN, edgeR, ComplexHeatmap, and clusterProfiler were used for specific analyses as described in the corresponding figures and result section.

---

**Supplementary Figure 1.**
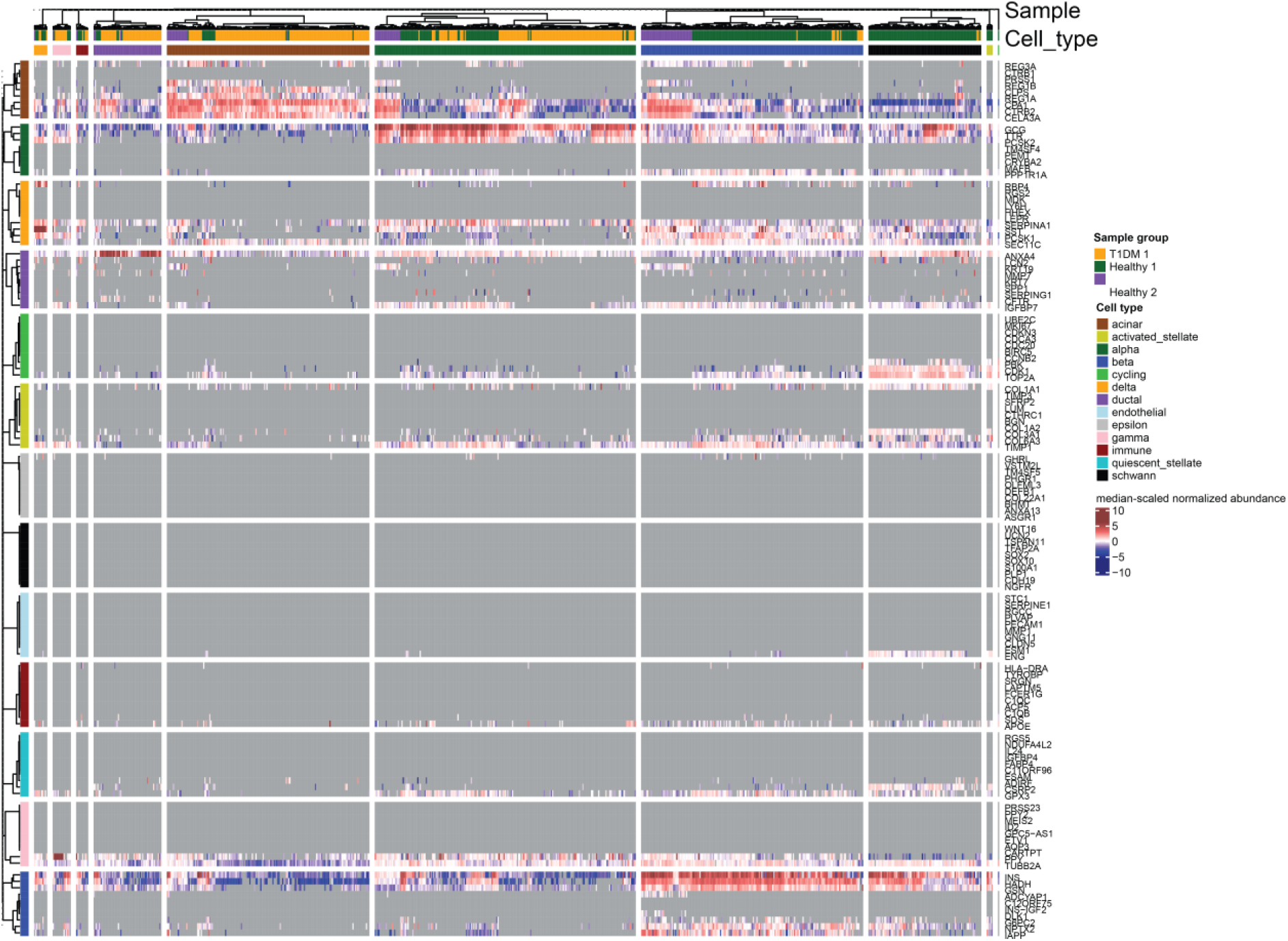
Expression of the anchor markers from Azimuth. Heatmap of the median-scaled normalized abundance of the corresponding protein to the specific anchor genes detected in the healthy 1, healthy 2, and T1DM 1 datasets. Grey colour on the heatmap represents missing values. Cell types were annotated using Azimuth as in Figure 2c.

**Supplementary Figure 2.**
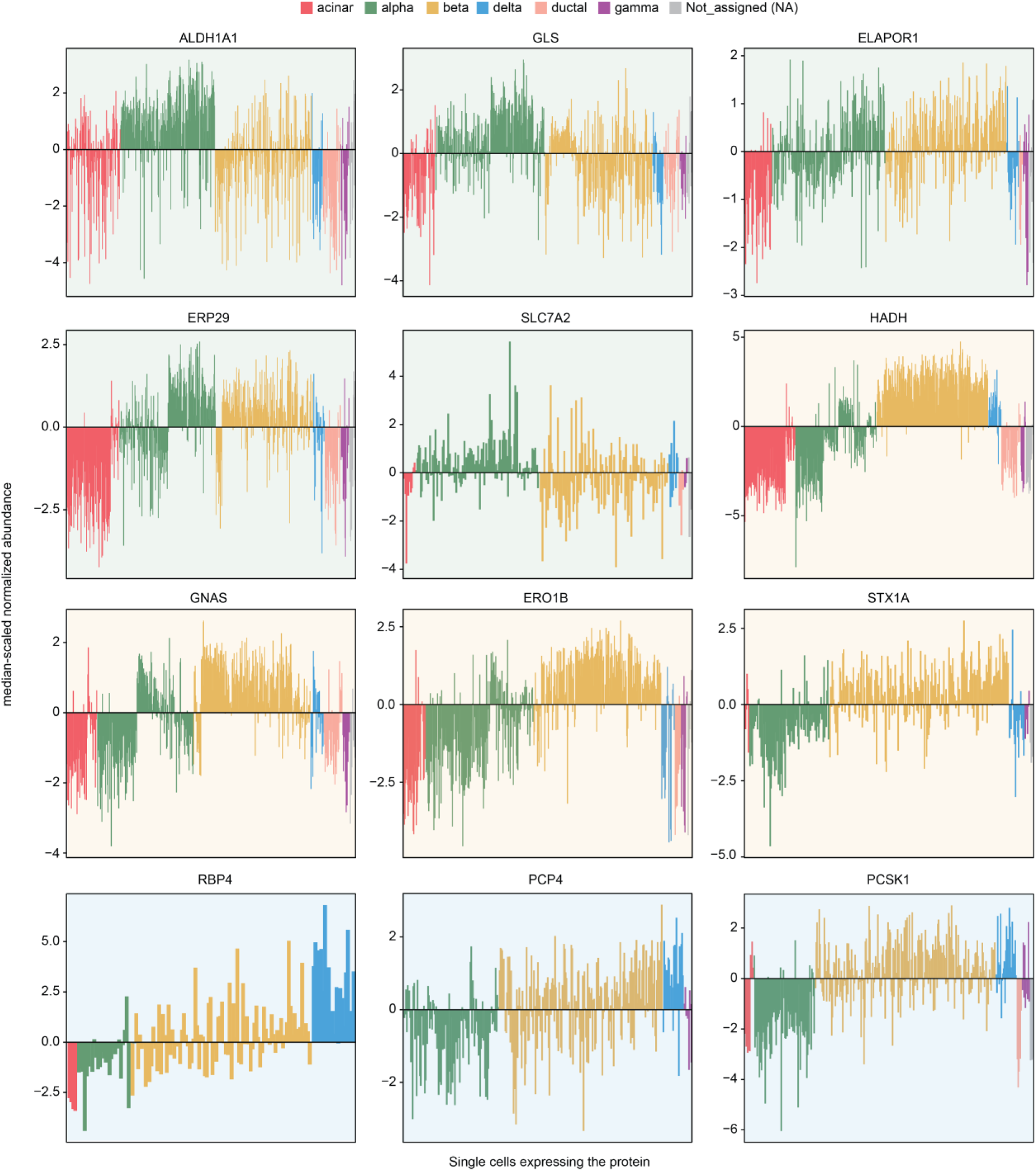
Relative abundance of proteins referenced as markers of alpha, beta, and delta cells. Median-scaled normalized abundance of proteins referenced as specific markers of alpha (green), beta (yellow), delta (blue) in Azimuth and in the only other SCP study of pancreatic islets from Fulcher et al., in each cell of the healthy 1, healthy 2 and T1DM 1 datasets. Only acinar, alpha, beta, delta, ductal, gamma and Not-assigned cells are shown. The annotated cell types are highlighted.

**Supplementary Figure 3.**
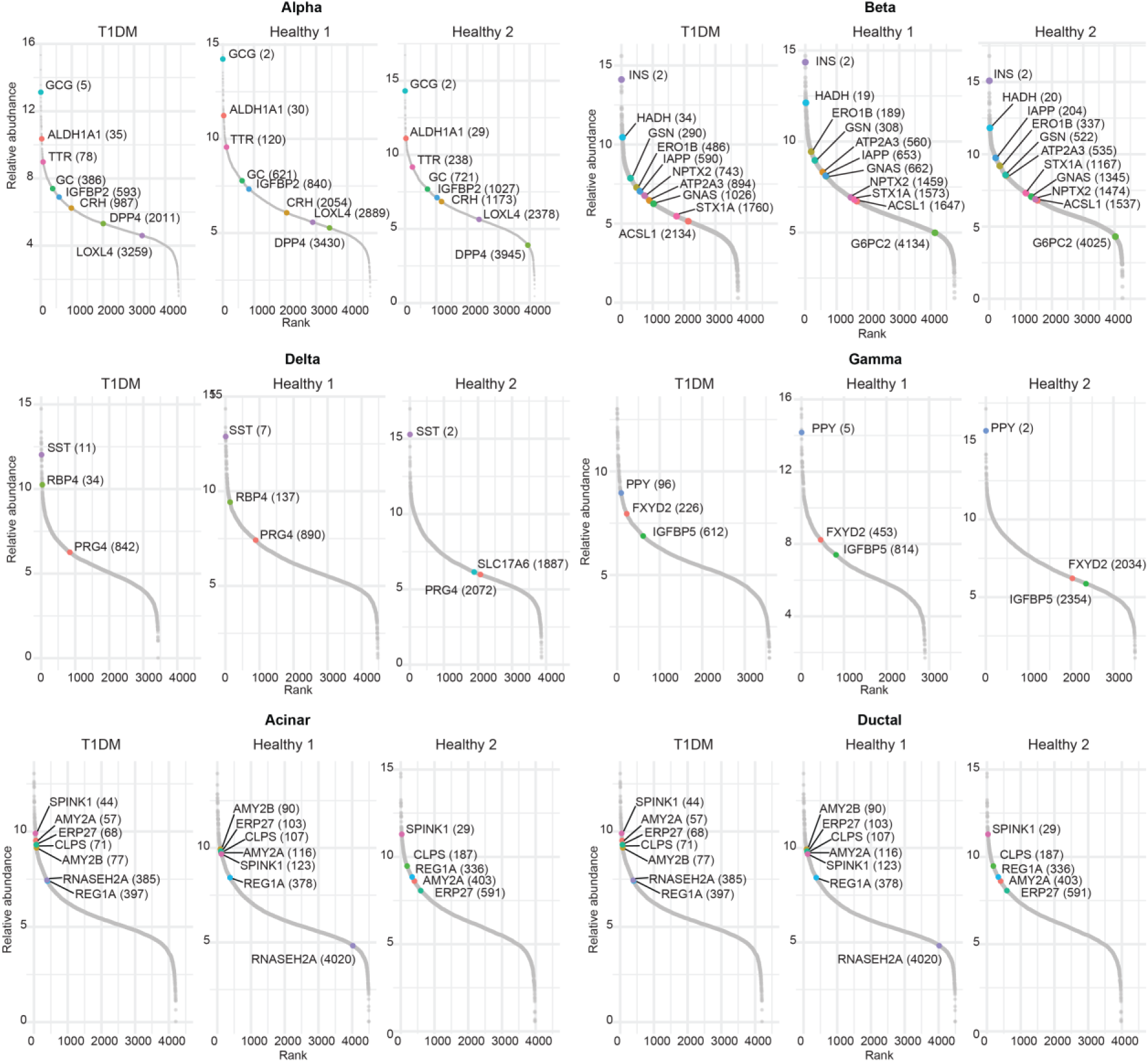
Ranking of the curated markers in the corresponding cell type. Rank (x-axis) according to the protein log2 LFQ intensity (y-axis) of proteins selected as markers for alpha, beta, delta, gamma, acinar and ductal cells.

**Supplementary Figure 4.**
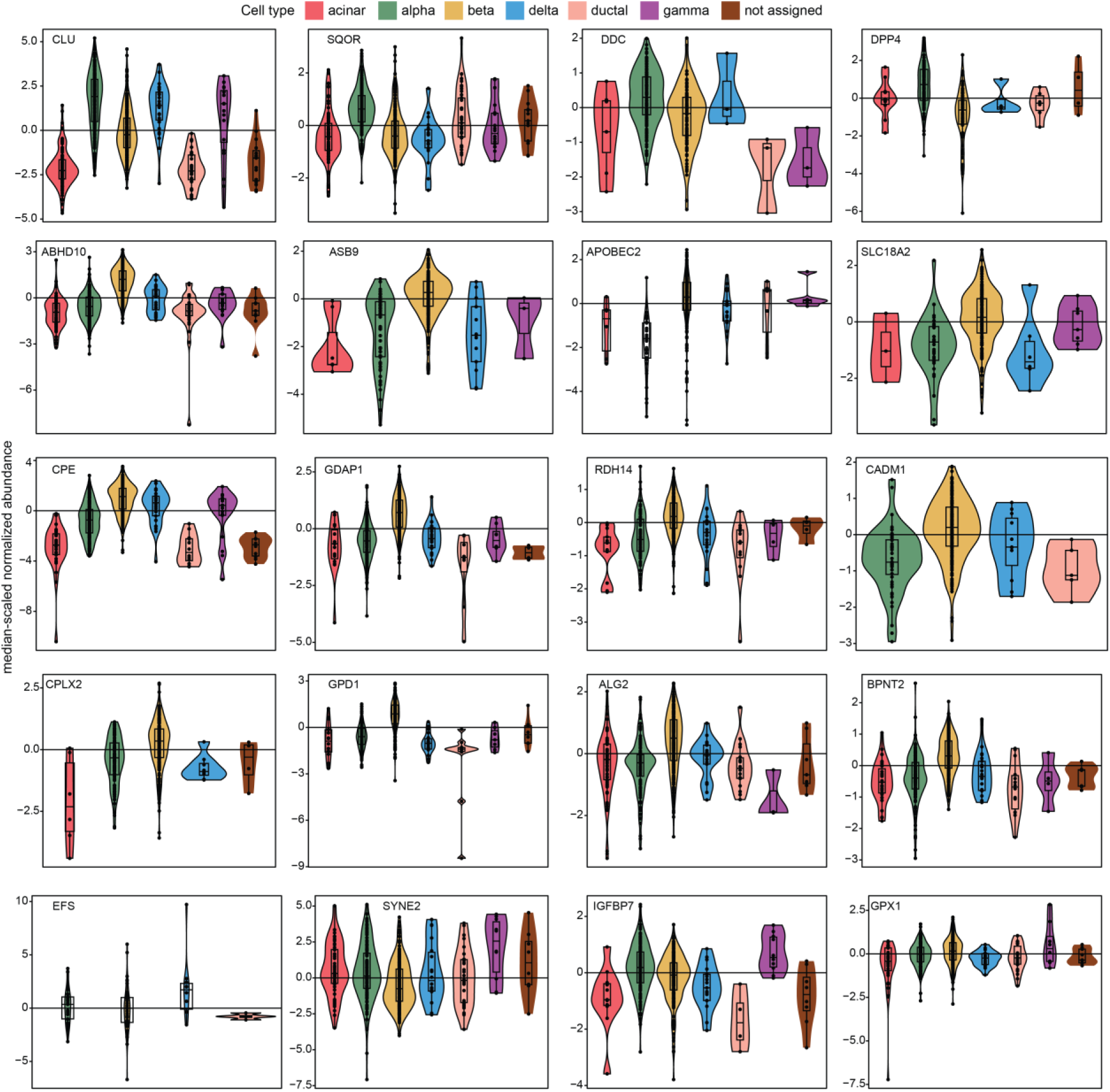
Abundance of differentially expressed proteins. Median-scaled normalized abundance of proteins showing specificity toward one cell type between alpha (green), beta (yellow), delta (blue) and gamma (purple) based on the differential expression analysis of Figure 5. The data encompasses healthy 1, healthy 2 and T1DM 1 datasets.

**Supplementary Figure 5.**
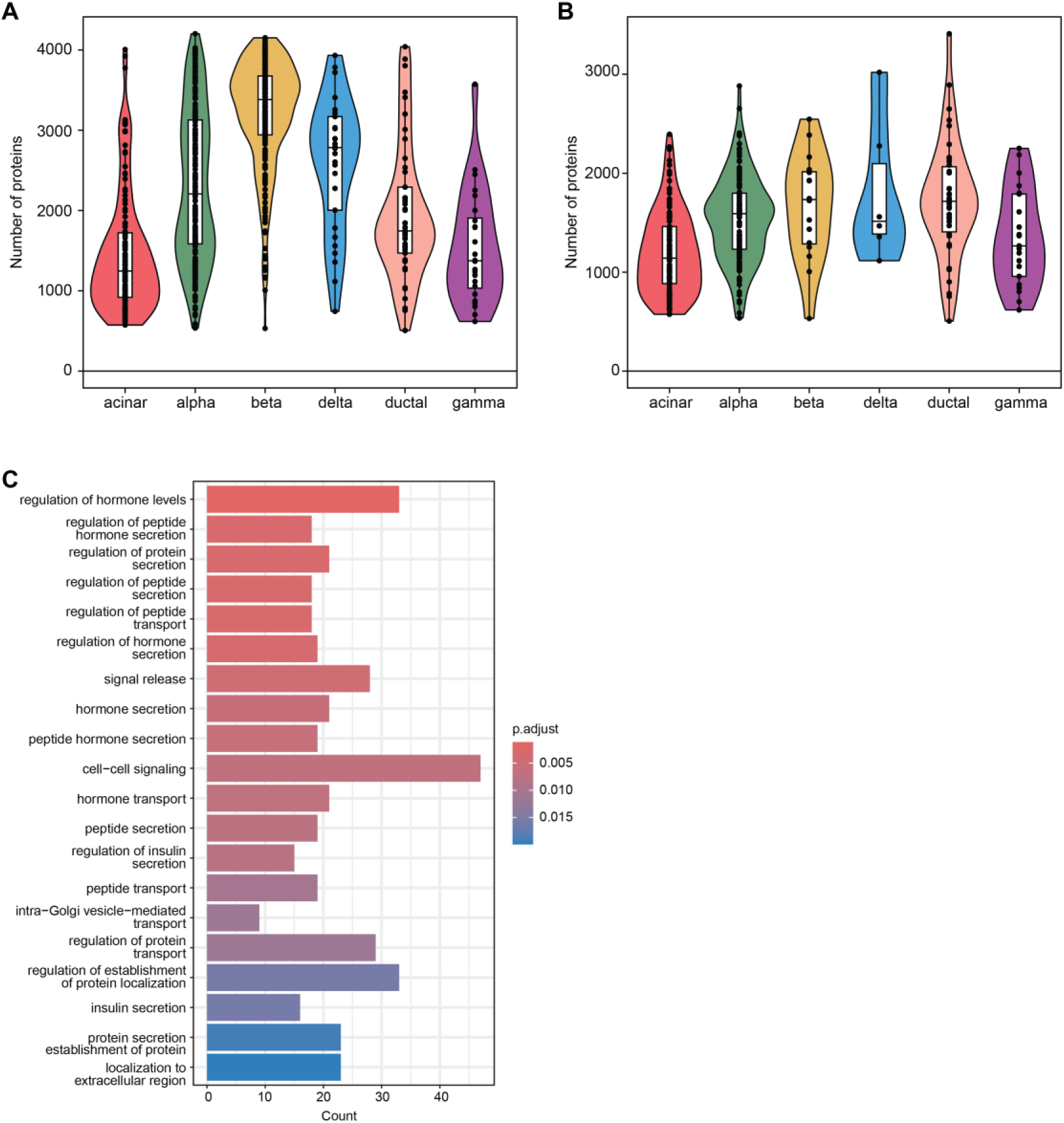
Number of proteins identified per cell type. (**A**) Number of proteins quantified in healthy 1, healthy 2 and T1DM 1 donors. (**B**) Number of proteins quantified in T1DM 1 donor. (**C**) Gene Ontology (GO) biological processes enrichment analysis using the clusterProfiler R package of proteins which are significantly differentially expressed in beta cells compared to other cell types (abs(log2 FC)>1 and adjusted p-value < 0.05). p-values were adjusted using the Benjamini-Hochberg procedure.

**Supplementary Figure 6.**
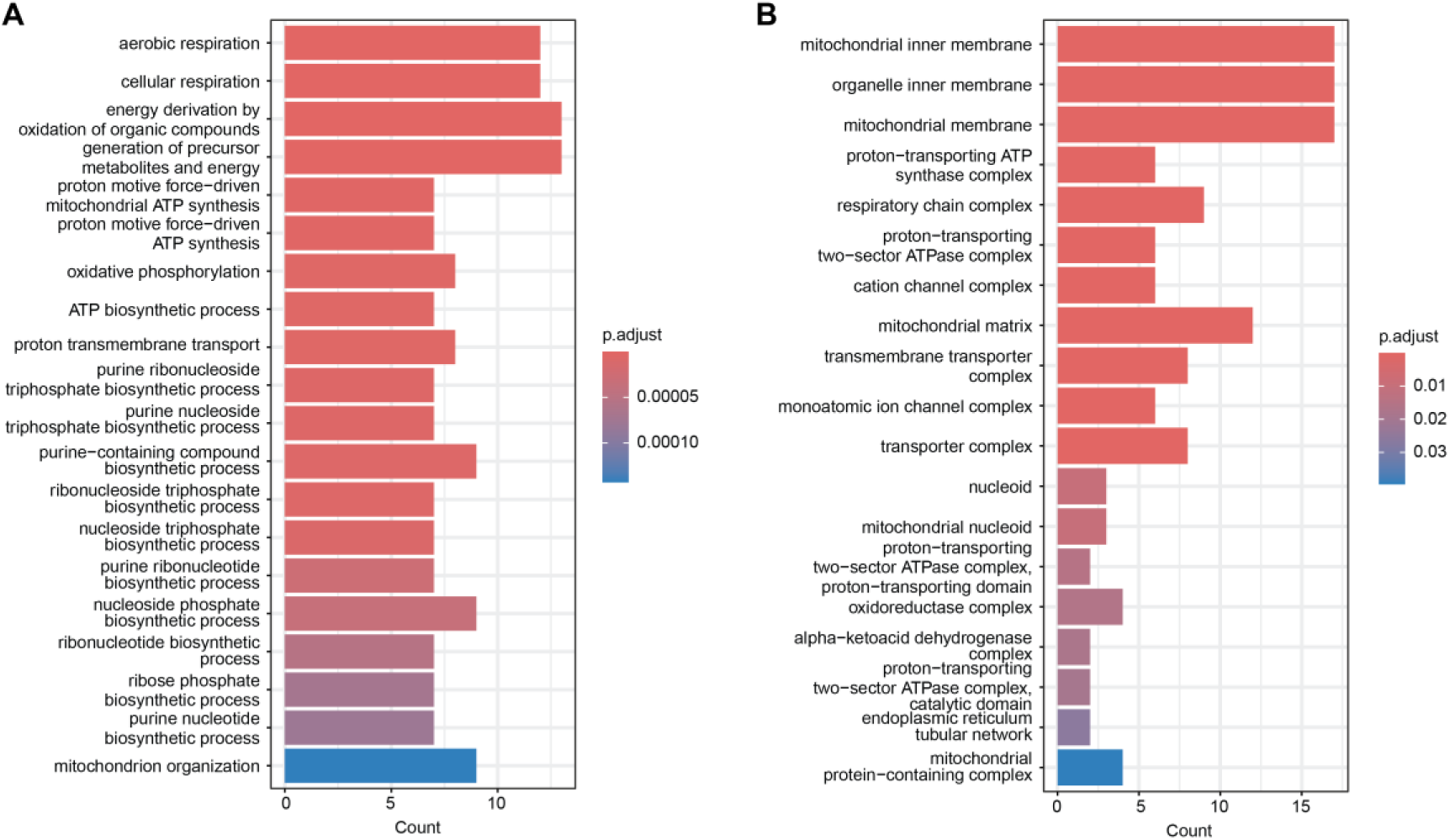
Gene Ontology (GO) enrichment analysis of proteins differentially expressed between alpha and beta cells in healthy 1 and healthy 2. (**A**) Enrichment of GO biological processes. (**B**) Enrichment of GO cellular compartments. The analysis was performed using clusterProfiler R package. P-values were adjusted using Benjamini-Hochberg procedure.

**Supplementary Figure 7.**
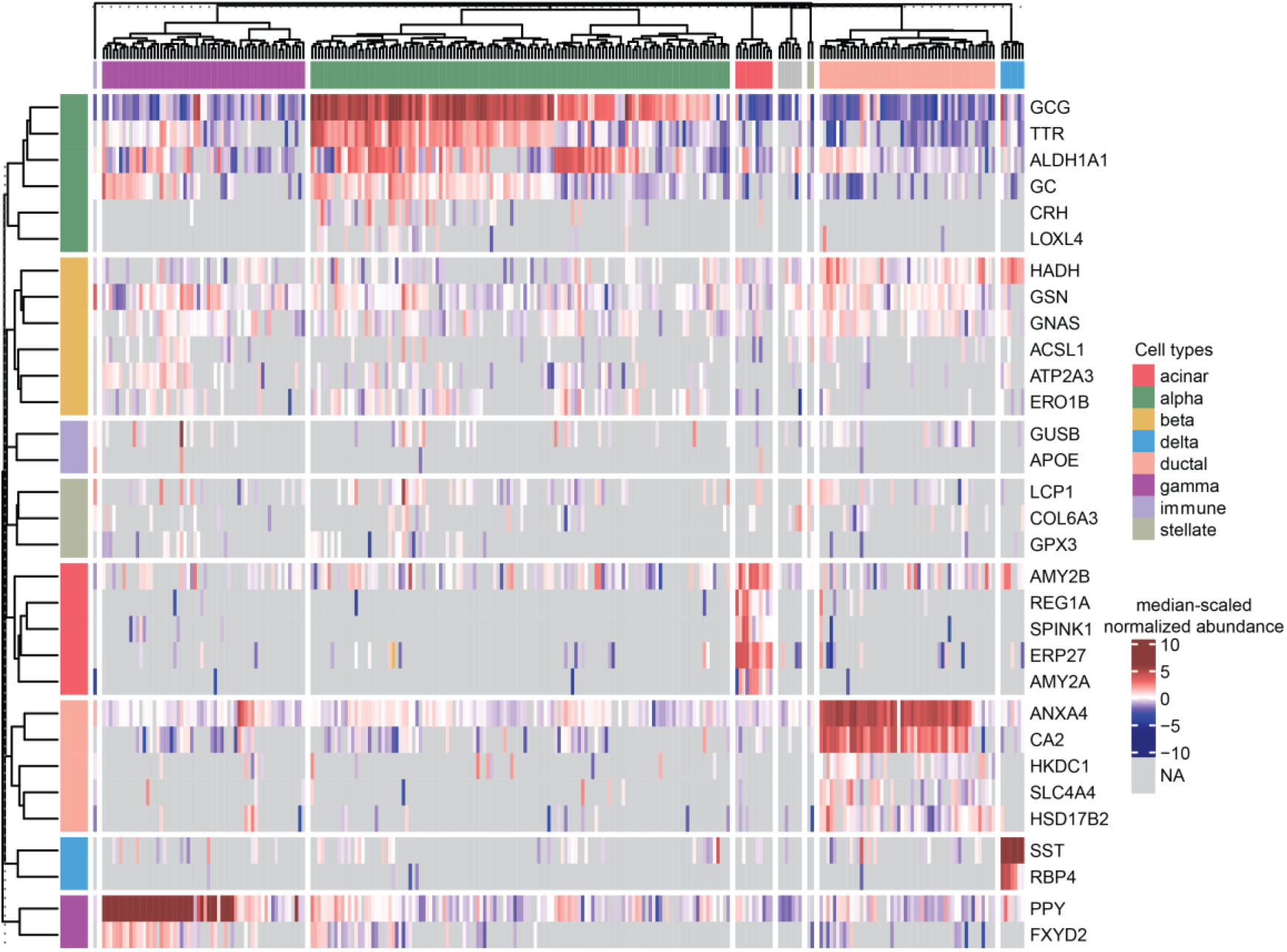
Abundance of markers in T1DM 2 donor. Heatmap of the median-scaled normalized abundance of the curated protein markers in the T1DM 2 donor dataset. Grey colour on the heatmap represents missing values. Cell types were annotated according to the same strategy as in Figure 3A.

**Supplementary Table 2.** Donor characteristics.

| Donor No. | Donor Type | Age | Sex | HbA1c (mmol/mol) | BMI | Q-test (SI) | Purity (%) | Isolation Date | Single-cell Production Date |
| --- | --- | --- | --- | --- | --- | --- | --- | --- | --- |
| Healthy 1 | ND | 44 | M | 37 | 35.6 | 13.7 | 90 | 240515 | 240521 |
| Healthy 2 | ND | 40 | F | 35 | 30.2 | 15.5 | 89 | 240617 | 240620 |
| Healthy 3 | ND | 50 | F | 48 | 35.4 | 7.8 | 96 | 240229 | 240305 |
| T1DM 1 | T1D | 42 | M | 49 | NA | 1.8 | 63 | 240421 | 240423 |

**Supplementary Table 3.**
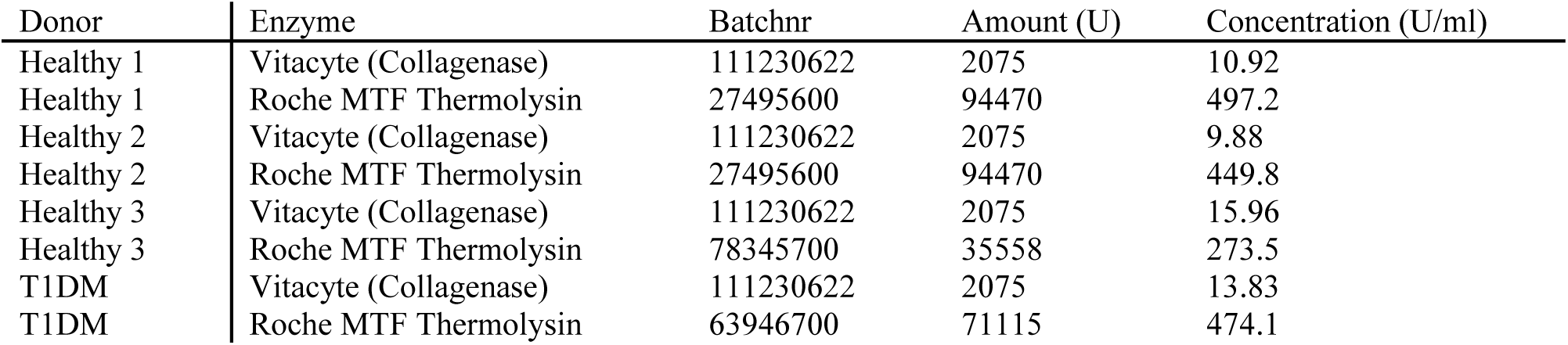
Enzymes and concentrations used for each donor.

| Donor | Enzyme | Batchnr | Amount (U) | Concentration (U/ml) |
| --- | --- | --- | --- | --- |
| Healthy 1 | Vitacyte (Collagenase) | 111230622 | 2075 | 10.92 |
| Healthy 1 | Roche MTF Thermolysin | 27495600 | 94470 | 497.2 |
| Healthy 2 | Vitacyte (Collagenase) | 111230622 | 2075 | 9.88 |
| Healthy 2 | Roche MTF Thermolysin | 27495600 | 94470 | 449.8 |
| Healthy 3 | Vitacyte (Collagenase) | 111230622 | 2075 | 15.96 |
| Healthy 3 | Roche MTF Thermolysin | 78345700 | 35558 | 273.5 |
| T1DM | Vitacyte (Collagenase) | 111230622 | 2075 | 13.83 |
| T1DM | Roche MTF Thermolysin | 63946700 | 71115 | 474.1 |

**Supplementary Table 4.** Antibodies and stain used for immunofluorescence staining.

| Antibody | Catalog # | Company | Concentration | Incubation time |
| --- | --- | --- | --- | --- |
| **Insulin** | 565689 | BD Biosciences | 1:100 | 1 h |
| **Glucagon** | 565891 | BD Biosciences | 1:100 | 1 h |
| **Somatostatin** | 566032 | BD Biosciences | 1:100 | 1 h |
| **Sytox Orange** | S11368 | Life Technologies | 500 nM | 5 min |

